# Leveraging artificial intelligence and software engineering methods in epidemiology for the co-creation of decision-support tools based on mechanistic models

**DOI:** 10.1101/2023.09.03.555060

**Authors:** Sébastien Picault, Guita Niang, Vianney Sicard, Baptiste Sorin-Dupont, Sébastien Assié, Pauline Ezanno

**Affiliations:** Oniris, INRAE, BIOEPAR, 44300, Nantes, France

**Keywords:** Epidemiological modelling, decision-support tools, intervention ranking, knowledge representation, domain-specific language, code generation

## Abstract

Epidemiological modelling is a key lever for infectious disease control and prevention on farms. It makes it possible to understand the spread of pathogens, but also to compare intervention scenarios even in counterfactual situations. However, the actual capability of decision makers to use mechanistic models to support timely interventions is limited. This study demonstrates how artificial intelligence (AI) techniques can make mechanistic epidemiological models more accessible to farmers and veterinarians, and how to transform such models into user-friendly decision-support tools (DST). By leveraging knowledge representation methods, such as the textual formalization of model components through a domain-specific language (DSL), the co-design of mechanistic models and decision-support tools becomes more efficient and collaborative. This facilitates the integration of explicit expert knowledge and practical insights into the modelling process. Furthermore, the utilization of AI and software engineering enables the automation of web application generation based on existing mechanistic models. This automation simplifies the development of DST, as tool designers can focus on identifying users’ needs and specifying expected features and meaningful presentations of outcomes, instead of wasting time in writing code to wrap models into web apps.

To illustrate the practical application of this approach, we consider the example of Bovine Respiratory Disease (BRD), a tough challenge in fattening farms where young beef bulls often develop BRD shortly after being allocated into pens. BRD is a multi-factorial, multi-pathogen disease that is difficult to anticipate and control, often resulting in the massive use of antimicrobials to mitigate its impact on animal health, welfare, and economic losses. The decision-support tool developed from an existing mechanistic BRD model empowers users, including farmers and veterinarians, to customize scenarios based on their specific farm conditions. It enables them to anticipate the effects of various pathogens, compare the epidemiological and economic outcomes associated with different farming practices, and decide how to balance the reduction of disease impact and the reduction of antimicrobial usage (AMU).

The generic method presented in this article illustrates the potential of artificial intelligence (AI) and software engineering methods to enhance the co-creation of decision-support tools based on mechanistic models in veterinary epidemiology. The corresponding pipeline is distributed as an open-source software. By leveraging these advancements, this research aims to bridge the gap between theoretical models and the practical usage of their outcomes on the field.

**Highlights:**

- AI make mechanistic epidemiological models usable to support decisions on farms
- Textual knowledge representation fosters co-design of mechanistic models and tools
- AI and software engineering automate web app generation to ease disease control

## 1 Introduction

Mechanistic modelling has been broadly used in human, animal, and plant epidemiology, especially in the two last decades. It helps understand and predict the spread of infectious diseases, based either on mathematical (Hethcote, 2000; Keeling, 2005; Keeling and Rohani, 2008) or agent-based methods (Amouroux et al., 2008; Bigley Dunham, 2005; Roche et al., 2008). Mechanistic epidemiological models are also highly valuable to assess the efficacy of control measures or compare intervention scenarios (Ferguson et al., 2005; Halloran et al., 2008; Thulke, 2011). They became gradually more and more used in veterinarian epidemiology to address epidemic livestock outbreaks, e.g. foot and mouth disease (FMD) (Webb et al., 2017). But, they also proved a powerful approach to manage endemic diseases, which have a lasting impact on the livestock sector, such as bovine paratuberculosis (Beaunée et al., 2017; Bennett et al., 2010) or bovine viral diarrhoea (Thulke et al., 2018). Indeed, they can represent concrete on-farm intervention levers and account for epidemiological, but also social, economic or environmental considerations. This makes mechanistic models highly valuable to build up recommendations for decision makers (farmers, veterinarians, public health managers, etc.) (Ezanno et al., 2020).

The development of mechanistic models for decision support has also increased drastically during the COVID-19 pandemic, leading to a multiplicity of open-source software tools (Danon et al., 2021; Hinch et al., 2021; Kerr et al., 2021; Reguly et al., 2022; Schneider et al., 2020). But, this also highlighted the difficulty to design, calibrate, validate, coordinate such models in a timely and transparent manner during major crises and use them for effective communication with decision makers (Barton et al., 2020; Meredith et al., 2021; Shea et al., 2023).

Hence, non-trivial mechanistic models are in general difficult to use by decision makers in an autonomous way. Indeed, epidemiological models are mainly developed in academic contexts, focusing on providing insights on complex research questions. They aim at simplifying field situations to capture the core dynamics of underlying processes: hence, they try to stay as simple as possible, involving mathematical approaches with as few parameters as required, to reproduce typical situations rather than a single specific observation. This sometimes also requires introducing finer-grained processes and parameters that are useful to represent the mechanisms involved in the system (e.g., pathogen characteristics), but may be not directly observable in the real system or very far from the stakeholders’ practical concerns. Parameters play multiple roles, e.g., shaping the behaviour of mathematical or rule-based processes involved in the core of the model, but also defining on/off switches for studying contrasted scenarios. The outputs of an academic model are composed of observables of interest with regards to the research questions and of additional variables that help characterize the behaviour of the model a posteriori.

To make such academic models evolve towards field applications, complementary processes must be taken into account, coming with more parameters. Such comprehensive models are more in line with expectations on the field, as decision makers expect to be asked for precise, concrete, and limited input information and provided with visual, unambiguous, and targeted output information, within tools that are sufficiently realistic for their conclusions to be meaningful in terms of farming and veterinary practices. But, as stated in (Divya et al., 2021): “comprehensive models may not be practical in field situations if they require inputs that are not easy to measure. The complexity of these comprehensive models makes them difficult to understand and more difficult to use and apply”. Especially, the visualization of model results is particularly important (Dykes et al., 2022; Muellner et al., 2018). Besides, to be used as relevant decision support tools, academic models need to be extended to account for a broader context, such as socio-economic issues (Sutherland and Freckleton, 2012).

Actually, in the majority of the work cited so far, academic models are developed first, and then, some of their results are expected to guide interventions. To benefit from the capability of mechanistic models to address a broad range of research questions, the ideal approach would consist in building tools on the top of existing models, targeting a specific practical issue for a dedicated category of users. This is feasible and has been done out on a case-by-case basis (Ezanno et al., 2020), but requires a long and tedious development effort to write the application that is able to interact properly with the underlying model and produce relevant and intelligible results. Indeed, models aimed to be useful for veterinary medicine involve several processes in addition to infection to accurately account for population dynamics and management, observable clinical signs, disease detection methods (visual appraisal, individual or collective tests, etc.), and the multiple interventions that can be implemented to mitigate disease impact. Thus they result in complex simulation codes which are difficult to write, check, maintain, and share with non-modellers, impeding both reproducibility and confidence in the models, which are seen as black boxes (Peng, 2011). Besides, updating the tool or the model requires to modify the whole software.

To overcome those limitations, we developed a method for automating the production of web applications based on existing mechanistic epidemiological models. Such applications can be co-designed with end-users such as veterinarians, farmers, technical advisors in the livestock sector, or public health services, to identify their practical concerns and formalize them in a structured text document. This text file is then automatically processed to produce the code for a web app which simulates and compare the users’ scenarios of interest and provides them with meaningful outputs. This work relies upon a combination of AI methods (knowledge representation, agent-based architecture) and software engineering methods (domain-specific languages), both becoming increasingly important in animal health research (Ezanno et al., 2021).

## 2. Materials and methods

### 2.1 Leveraging AI for mechanistic epidemiological models

In this study, we used the EMULSION framework (Picault et al., 2019) to develop human-readable mechanistic epidemiological models while reducing development time. EMULSION is an open-source software, distributed as a Python module (on the PyPI repository and from https://sourcesup.renater.fr/www/emulsion-public). This generic approach is based on the combination of two AI methods.

First, the components of an epidemiological model (model structure, processes, epidemiological units, time scale, parameters, mathematical functions, initial conditions, observable variables…) are described as a structured text based on the YAML syntax (Ben-Kiki et al., 2009) which looks like a bullet-point document (mainly based on lists and key-value pairs). Indeed, it is crucial to keep the model human-readable as long as possible, because as soon as a model is implemented into computer code, its capability to be checked, discussed, revised, maintained but also turned into a DST is drastically reduced. Knowledge representation approaches, coming from symbolic AI (Russell and Norvig, 2010), make it possible to overcome this issue by providing methods to process explicit, human-readable descriptions of the content of mechanistic models and of their outcomes. EMULSION relies upon a domain-specific language (DSL) to formalize models. Contrary to a general-purpose programming language which can produce any kind of computation, a DSL is aimed at expressing the knowledge or processes attached to a reduced application field. DSLs are at the crossroads between AI and software engineering and proved a particularly efficient way to implement a “low-code” approach (Bucchiarone et al., 2021) that facilitates interactions between all scientists involved in the study of a specific pathosystem.

Second, EMULSION models are read by a generic simulation engine to process the simulation. To do so, the simulation engine has to be able to cope with a diversity of modelling methods (individual-based vs. compartment-based, with dynamics driven by equations, probabilistic behaviours, decision rules, etc.) and scales (individuals, groups, populations, metapopulations). In the field of AI, the recent development of multi-level agent-based architectures (Mathieu et al., 2018) emerged as a solution for representing heterogeneous entities at various scales, endowed with behaviours of their own. This approach was implemented in EMULSION so that the simulation engine could read the DSL, dynamically create agents representing the required granularity levels with relevant behaviours and process the simulation.

Since its first release (Picault et al., 2019), EMULSION has been extended to also account for complex population structuring (Sicard et al., 2021), which are quite common in veterinary epidemiology, such as the combination time, group and building constraints found in batch-rearing systems in the pig sector (Sicard et al., 2022a). This multi-level agent-based approach has been incorporated in the simulation engine and declarative DSL of EMULSION recently (Sicard et al., 2022b).

### 2.2 Web application architecture for model-based DST

The use of realistic mechanistic models in veterinary epidemiology may require a substantial computing power. Indeed, assessing for instance the effectiveness of eradication control measures, or accounting for the intrinsic variability of biological processes or for the small size of within-farm animal populations, lead to stochastic models. Thus, several stochastic repetitions of each scenario must be run to provide analysable results. Besides, using models to support decision also implies to compare many scenarios, e.g. a scenario with or without the disease, with or without interventions, etc. Thus, it is not reasonable to envisage DST based on mechanistic models as lightweight applications that could be installed by each end-user. Instead, web applications are an easy way of making information accessible, and they can be used from most platforms (desktop computers, laptops, mobile phones, under most operating system).

Hence, to support such DSTs, we designed an architecture based on the interactions between a calculation server on which mechanistic simulations are run and their outcomes processed, and a web server which hosts DST applications. End-users are expected to submit specific information on practical cases, e.g., the population structure in their farm, the nature of vaccines or tests used, or economic values such as costs and prices. This information is processed by the web application to request simulations with specific parameters from the calculation server, which runs the corresponding scenarios from the mechanistic model, processes the raw outputs and sends back results to the app, which finally presents the relevant graphical outcomes to the users (Fig. 1)

**Figure 1.**
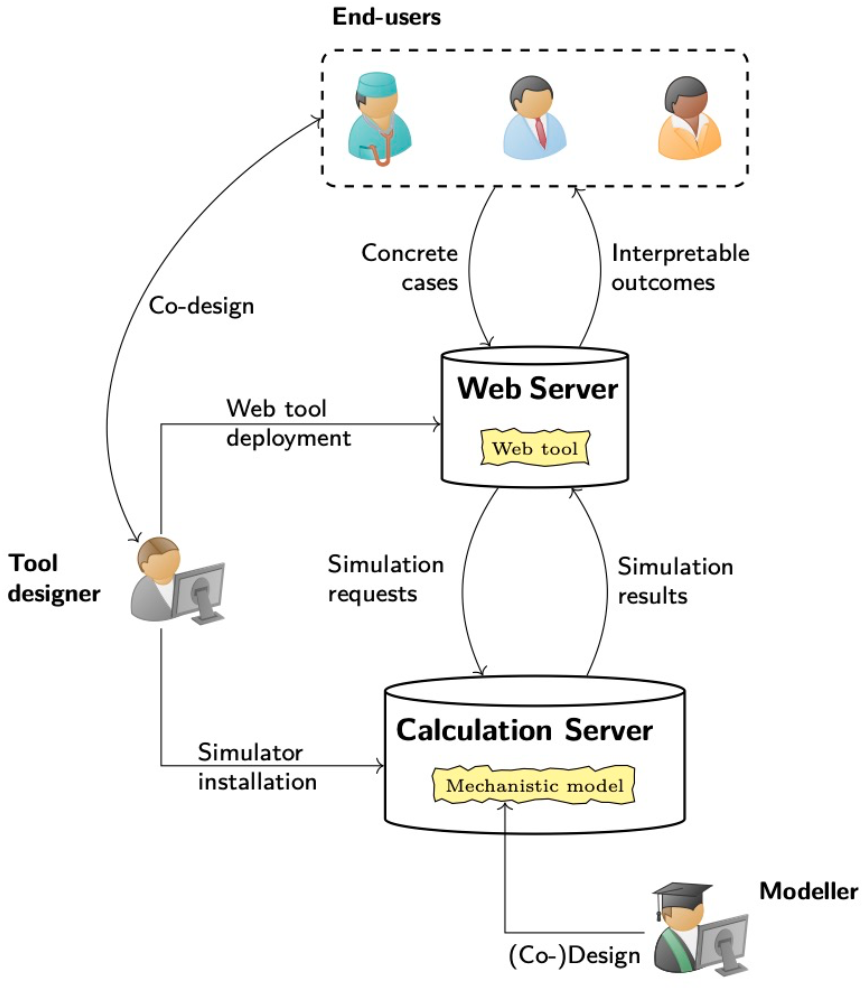
Architecture of a web decision-support tool based on a mechanistic model.

In that process, we assumed that the model had already been designed by a modeller, as much as possible in collaboration with potential or actual users (including other scientists). Three main tasks were left to the tool designer: first, identify DST features together with the end-users; second, produce the code of the web application from those specifications; and third, deploy the simulator on the calculation server and the web application on the web server. The deployment task should not raise issues but purely technical. As for the two others, we designed a declarative DSL to formalize the specifications of the web application in a human- and machine-readable format, and then a software engine to process them and automatically produce the code for the web application. The resulting framework, PASTE (Platform for Automated Simulation and Tools in Epidemiology), is described in the next section and summarised on Figure 2.

**Figure 2.**
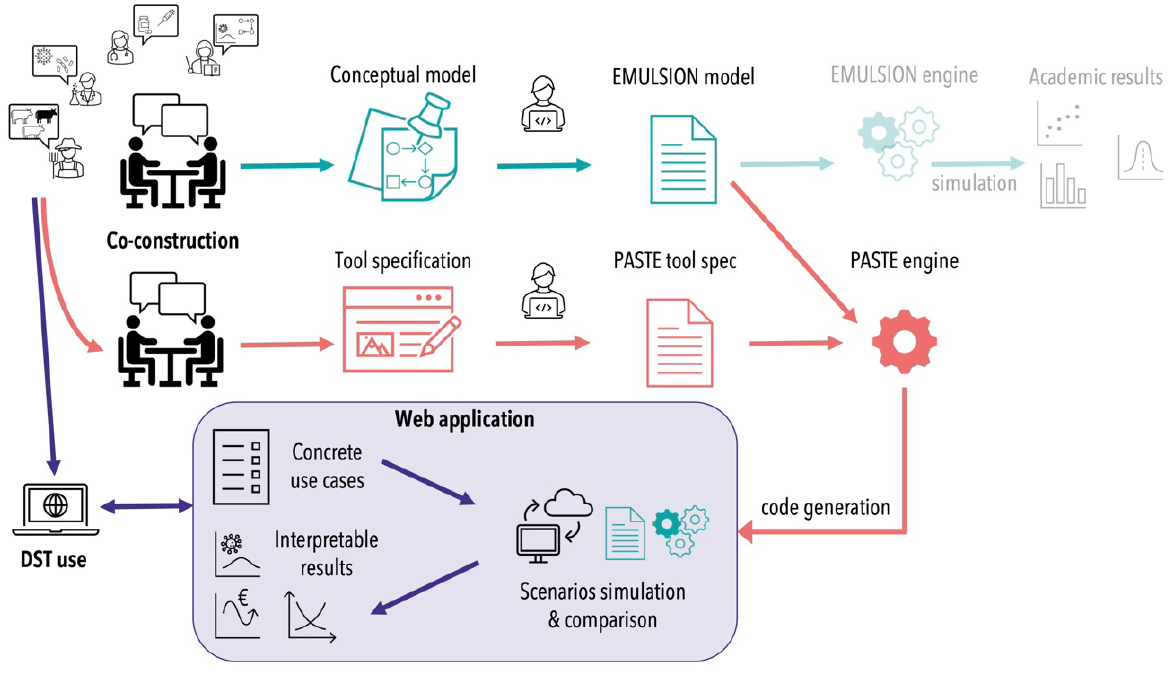
Production of a decision support tool as a web application by the PASTE software. (From an existing epidemiological model based on EMULSION). The key principle is to co-designed web tools as well as mechanistic models and formalize the result within a DSL. In EMULSION, the resulting model (a YAML document file) is usually processed for academic purpose; in PASTE, the tool specification (another YAML document) serves to generate the code for a web application, which in turn will use EMULSION engine to simulate the underlying EMULSION model with a specific setup and process the results to present what is relevant to the users’ concerns.

### 2.3 A declarative language for tool specification

As in EMULSION, the DSL designed for specifying the features of the web DST is based on YAML to foster easy reviewing and revisions in a declarative way. The DSL aims at representing several aspects of the model-based application: the link with the mechanistic model to use (written with EMULSION), the input forms and tool parameters to propose to users, the definition of the scenarios to compare (based on specific values for model parameters), the description of statistical treatments to perform on raw model outputs for each scenario and for scenario comparison, and finally the description of the graphical charts to display to the users.

In this DSL, input parameters can be freely set by the user, picked within predefined values or ranges, or defined as a group in an external CSV file. For instance, several pathogens can cause the same disease independently. To make the model pathogen-specific, many parameters have to be defined but then the parameter set should be kept consistent for this specific pathogen and not allow parameters to vary one by one. Hence, it is easier to group them in a CSV file where each line defines a set of parameter values associated with the name of a specific pathogen, so that users just have to choose a pathogen name. Scenarios are built from the combination of components corresponding to a concern (e.g. the initial prevalence, combined with the possible interventions), which are associated with specific parameter values in the model.

Due to the separation between simulation code and model description in EMULSION, the name, role and default value of parameters and outputs used by the mechanistic model can be easily distinguished from those that are introduced in the DST, either to pilot the execution of specific simulations on the server with the proper parameter values, or to transform model outputs into results that make practical sense for the end-users. For instance, the model may involve a parameter for the initial disease prevalence, whereas the tool may rather ask the user for an observed prevalence: then, the link between both can be made e.g. in the scenario definition, to calculate a true prevalence used by the simulations, based on additional knowledge on detection practices and their characteristics, such as the detection test sensitivity and specificity (collected as input parameters). Or, the tool may calculate financial losses or gains from parameters that are not present in the mechanistic model, such as the cost of veterinary interventions (visits, tests, vaccines, treatments…), the purchase and selling prices of animals, the feeding cost, etc. This can be achieved e.g., by using summary statistics based on the raw model outputs, such as the distribution of the number and type of interventions, the occurrence and duration of health issues, etc.

The specification of statistical treatments and graphs is inspired by the consecutive operations performed in the R library “dplyr”, and by the addition of graphical layers in “ggplot2”, respectively (Fig. 3). This helps tool designers who are already familiar with those libraries and is anyway close to classical request or graphic processing languages (e.g., SQL or D3JS). Examples of the PASTE DSL syntax are provided in the examples below (Fig. 5, Fig. 7, Fig. 9, and also throughout the Supporting Information).

**Figure 3.**
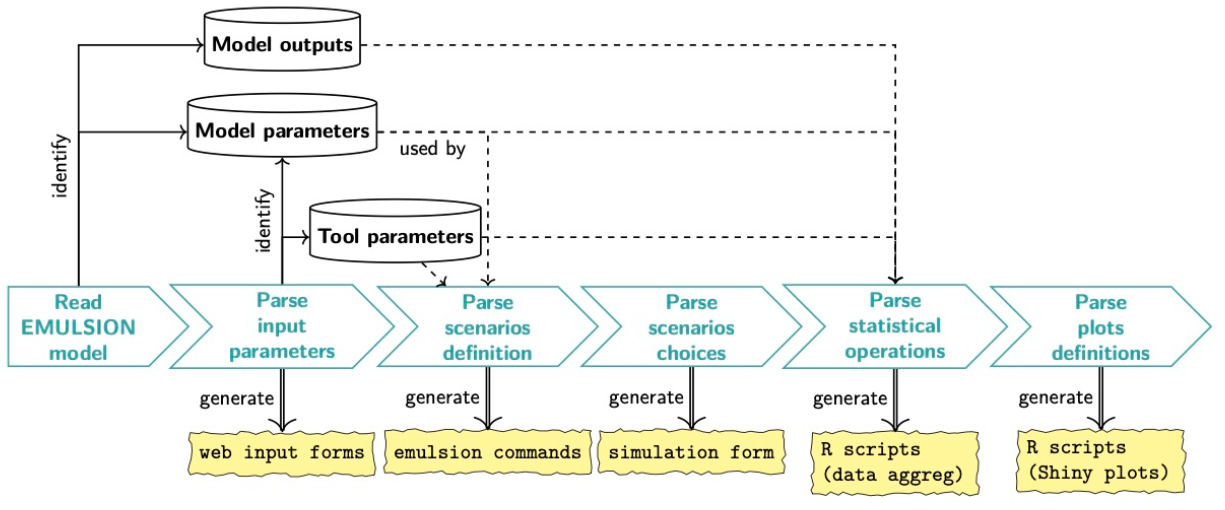
Overview of the successive steps to parse the tool specification file. After identifying the names of model parameters, model outputs and tool-specific parameters, the workflow consists in generating Python/Django code for input forms, preparing EMULSION commands with parameter values set according to the definition of scenarios, and R scripts to calculate relevant outputs and graphs for end-users from simulation outputs.

**Figure 4.**
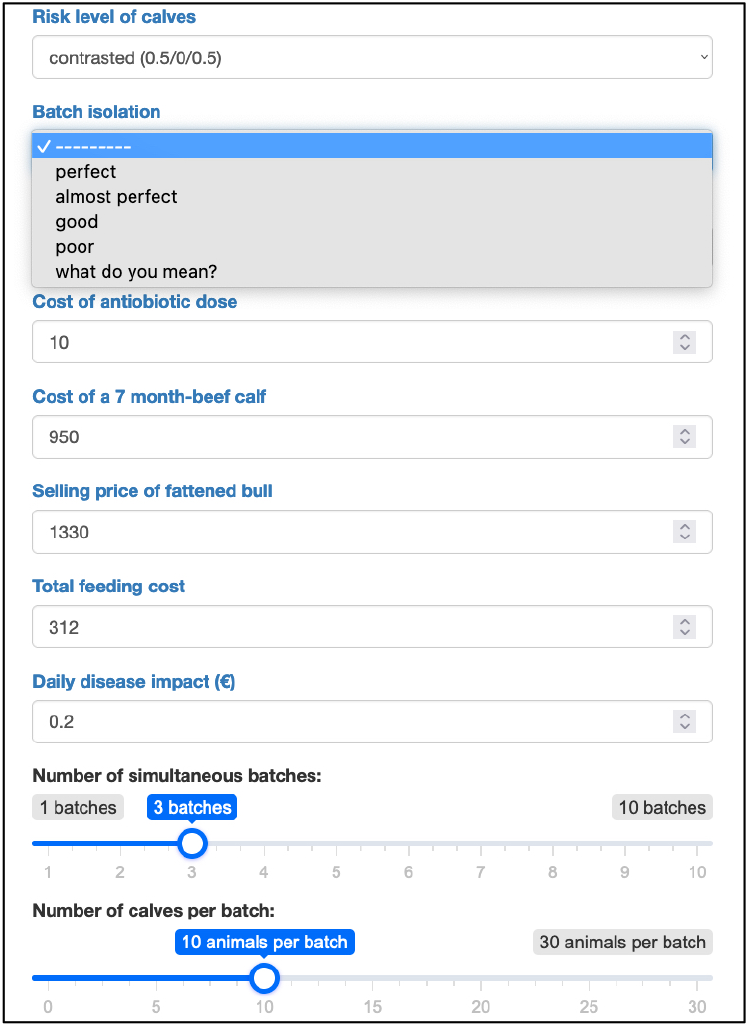
First input form. The user can indicate specific fattening conditions, such as the risk levels of beef calves in the batch, the level of isolation between batches, and the dominant pathogens (masked here by the drop-down menu), the composition of the farm and batches, but also costs and prices. The corresponding parameters are defined in the YAML file L30-46 and the input form itself L80-165 (Fig. 5).

**Figure 5.**
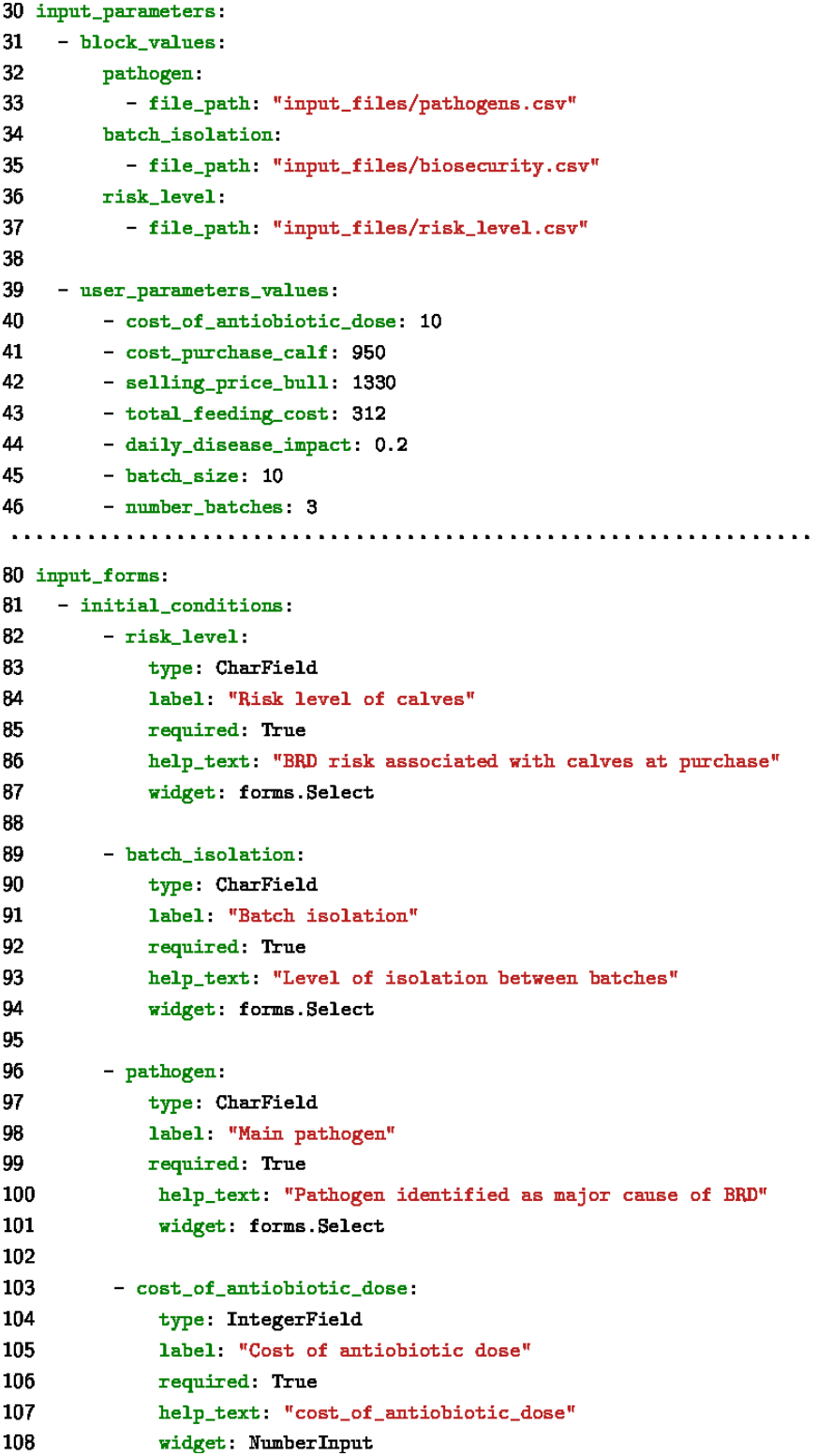
Specification of the input parameters and four first fields of the “initial conditions” input form (Fig. 4)

### 2.4 A software engine to generate the web application

The code architecture generated by the PASTE software engine is based on Django, a broadly used Python framework (https://www.djangoproject.com/), for managing the whole web application, and on R (https://www.r-project.org/) and common libraries (tidyverse, R-Shiny) to handle the statistical operations on model outcomes and the presentation of the graphical outputs.

The PASTE engine was designed to parse the YAML file containing the specification of the web application to build and generate the associated code, i.e. the Django objects and R scripts corresponding to DST components (Fig. 3). At each execution, this engine checks the validity of the statements with respect to the associated EMULSION model. First, the engine adapts existing templates and writes Python code where needed to create the input widgets (e.g., number inputs, sliders, checkboxes…) on the successive input forms (an example on a concrete application is provided on Figure 4 with the associated part of YAML specification on Figure 5). During that step, the PASTE engine also builds a representation of the parameters used in the tool and compares them to those defined in the EMULSION model.

Then, the engine examines the definition of scenarios. Scenario components are based on model or tool parameters. They can either re-use parameter values from given by the user in the input forms or default values defined in the model, or override those values to represent a specific situation. For instance, if an input form asks for the initial prevalence, a scenario component can leave this value unchanged to calculate a reference scenario, while another scenario component can automatically put the value to zero to represent a disease-free situation. Scenario components can also handle parameters which are not in the model, but just used in the user outputs (for instance, to explore high or low costs). The parsing of the scenario produces a web form where the users can choose what they want to compare (Fig. 6-7). It also produces a set of EMULSION commands, based on the proper parameter values, ready to be run on the calculation server when the tool will be used (Fig. SI 1).

**Figure 6.**
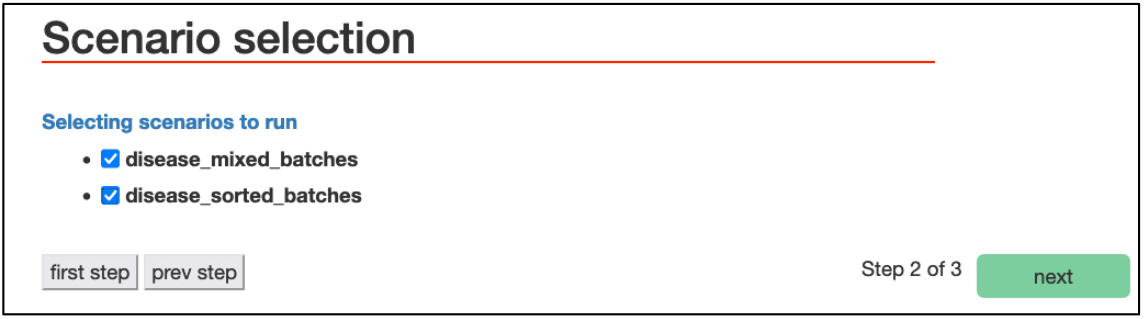
Form that proposes a choice on the scenarios to compare. Scenario components and their combination into actual scenarios are defined L50-78 (Fig. 7)

**Figure 7.**
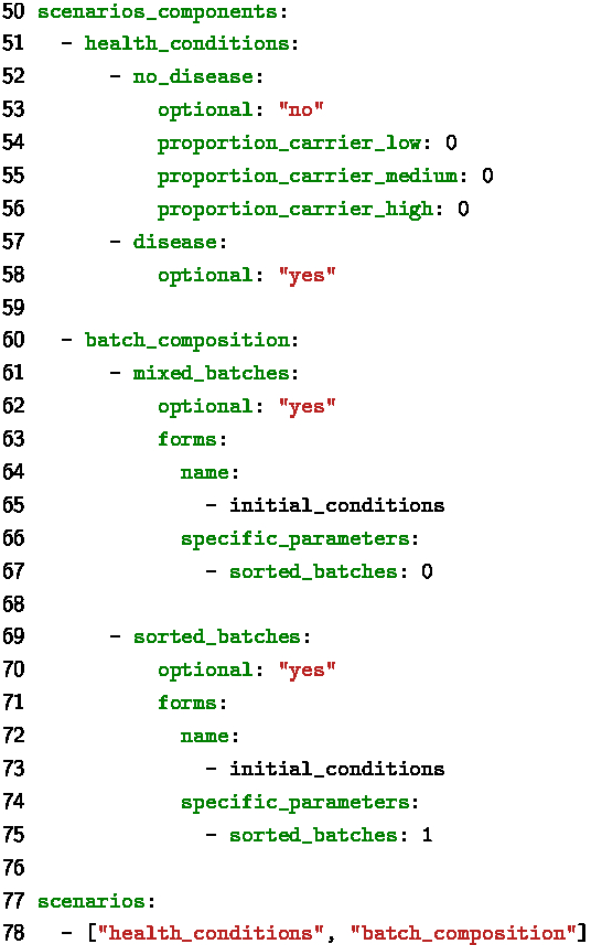
Specification of scenario components and scenarios (combination of scenario components). The scenario component “health_conditions” contains a part which is not optional (“no_disease”, L52-56): it will be run systematically to provide a baseline where no BRD cases occur (hence, not proposed to users in Fig. 6).

Third, the PASTE engine parses a section dedicated to statistics, to produce R code aimed at processing raw simulation results directly on the calculation server. Such treatments can be performed for each scenario (for instance an aggregation over the stochastic repetitions) or between some scenarios for comparison purpose. The specifications found in the YAML file are designed to be quite directly translated into R instructions, based on the “dplyr” library.

Finally, PASTE does the same for graphical results, based on the datasets produced by the statistical treatments. The plots can be distributed over several output pages or tabs to organize the results in a relevant way, and can either juxtapose or superpose scenarios (Fig. 8-9 and Fig. SI 3).

**Figure 8.**
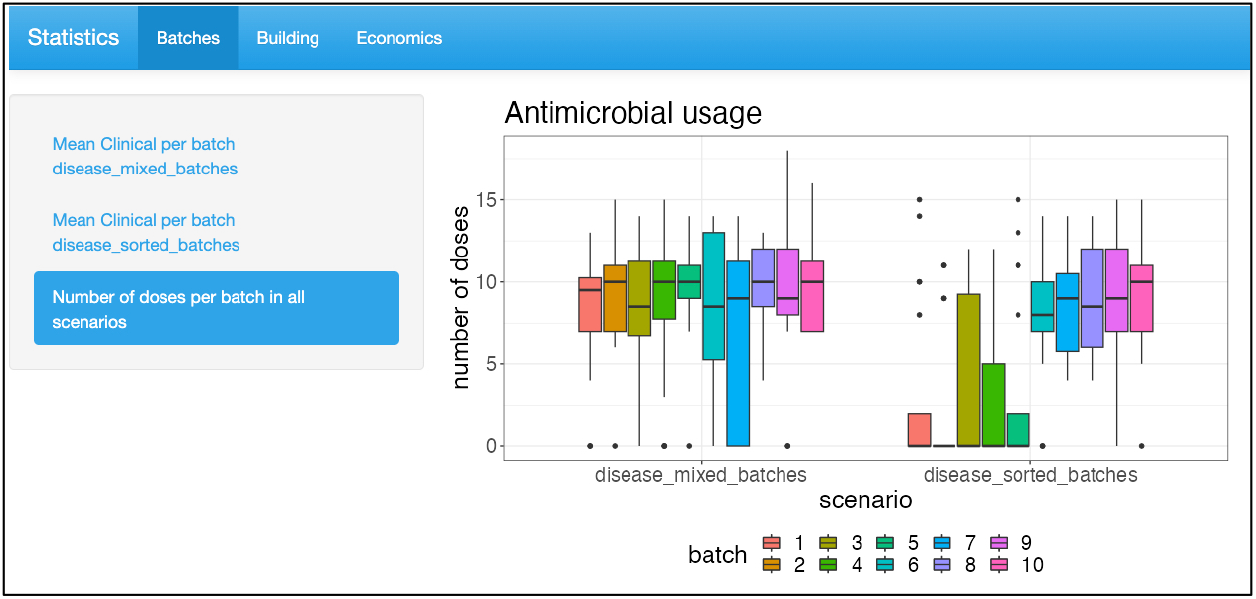
One of the output pages, showing results at batch scale. The figure presents the number of antimicrobial doses used in each batch (distribution over the stochastic repetitions) in two scenarios (left vs. right). The graph definition is L340-356, based on the treatment of simulation results specified in L213-220 (Fig. 9).

**Figure 9.**
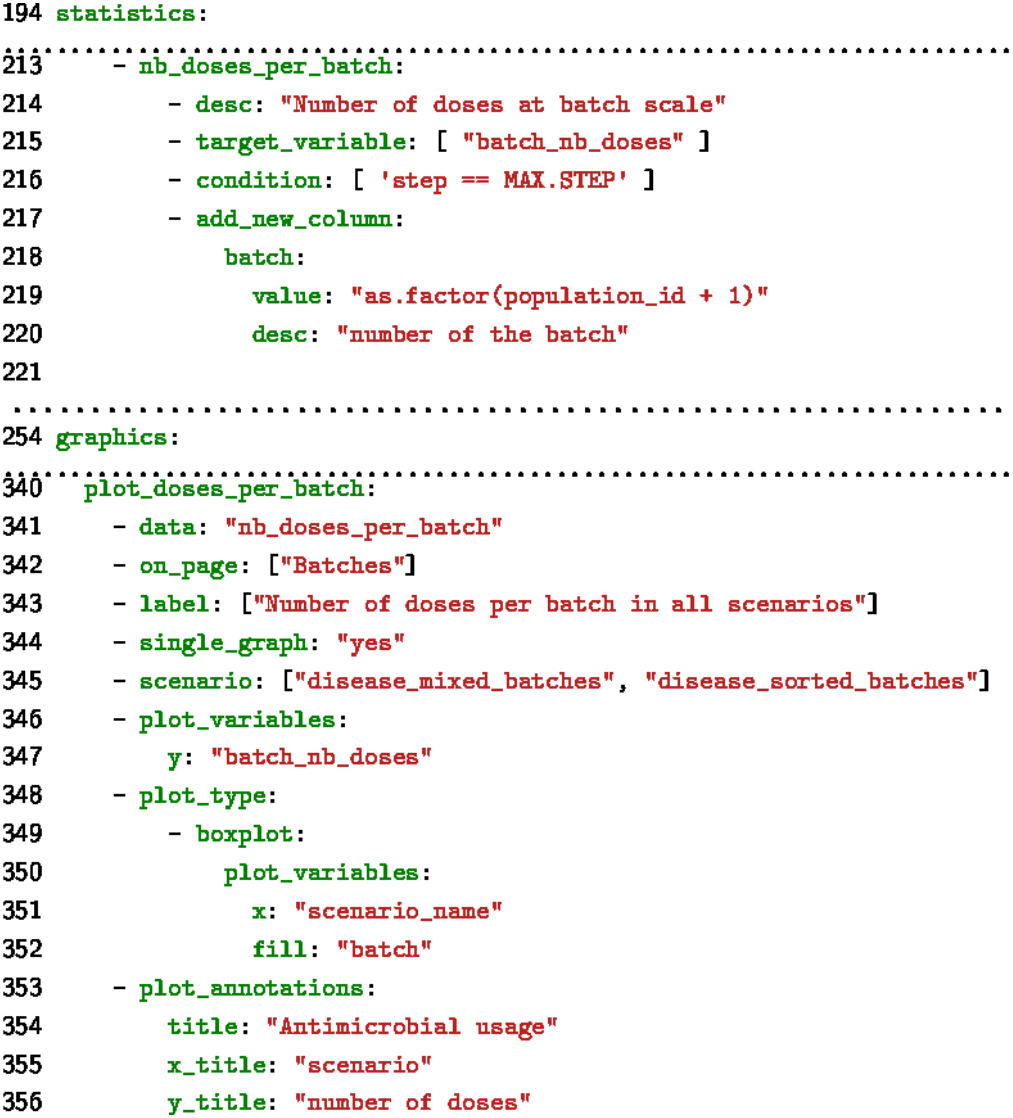
Specification for the summary statistics and the associated graphical representation (Fig. 8)

### 2.5 Proof of concept: an application to Bovine Respiratory Disease

To illustrate our approach, we used an existing model of Bovine Respiratory Disease (BRD) within a fattening building composed of multiple simultaneous batches (Sorin-Dupont et al., 2023)^1^. The model accounted for the within- and between-batch spread of pathogens, the onset of clinical signs, the detection of diseased calves and their treatment. Young beef calves arriving on farm after weaning were characterized by a risk level (to represent various risk factors such as distance travelled, mixing between animal sources, etc. in an aggregated way). Initial conditions allowed either to mix animals from different risk levels within each batch, or to allocate animals with the same risk level to the same batch. Finally, the infection process was designed, and the related model parameters calibrated, to account for three contrasted pathogens involved in BRD: the Bovine Respiratory Syncytial Virus (BRSV), *Mannheimia haemolytica*, and *Mycoplasma bovis*. The model was implemented with EMULSION 1.2.

The Decision-Support Tool we built on top of this model was intended to help fattening farmers assess the added value or risks associated with mixing young beef bulls, whatever their individual risk level of developing BRD. This indeed reflects current practices, as it is simpler to calculate fattening portions for batches of animals of similar weight, even coming from many distant farms. The proposed alternative was to allocate animals into batches based on their risk level.

To make the results more meaningful to farmers, treatment costs and animal prices, as well as disease impact (reduction of the weight intake) were introduced to calculate the gain of each management strategy. We defined four distributions of risk levels by defining four sets of parameter values in a CSV file (and summarized as “mainly low”, “mainly high”, “balanced”, “contrasted”). Similarly, we grouped parameter values associated with each pathogen. Finally, we also defined five graduated levels of batch isolation (representing biosecurity measures to limit between-batch transmission).

The outputs were designed to present results at the batch and at the building scales. Screenshots of the resulting tool are presented in section 3.2 (and Fig. SI 1 to Fig. SI 8), and the whole YAML file containing the specification of the tool is provided in the Supporting Information.

## 3. Results

### 3.1 Software architecture and code availability

The PASTE software engine has been tested to automate the production of web applications built on top of existing EMULSION mechanistic epidemiological models and aimed at supporting veterinary decision (see the BRD app example in the next section). PASTE is now distributed publicly as open-source software, under the Apache 2.0 licence. It can be installed from the following Git repository: https://forgemia.inrae.fr/dynamo/software/PASTE-public-release (which also contains the user guide and concrete examples of decision-support tools based on several EMULSION models).

The web architecture generated from the YAML specifications by the PASTE engine is fully operational and easy to deploy on calculation and web servers. Once the YAML specifications have been written, the generation of the web code takes only 2-3 minutes. Then, the deployment is twofold: the web architecture code (Django application and R-Shiny server) must be installed on the web server side, while the model, the EMULSION simulation engine, and the R software, must be installed on the calculation server side, as well as a task manager (currently, the Python package Celery) to manage the parallelization of simulations and scheduling of statistical treatments.

The web applications produced by the EMULSION-PASTE pipeline should rather be considered beta-versions of DST. On the one hand, they provide a simplified and meaningful interface to an existing mechanistic model, but on the other hand, they may lack “professional” features, such as an institutional or corporate branding, a user authentication module, etc. Besides, though the tool specifications are easy to co-design with stakeholders, it is also very important to evaluate the ergonomics and relevance of the tool together with a panel of representative users. Their feedback is necessary to ensure the significance and usefulness of the tools, and it is easy to incorporate changes in the web application features and rebuild it in a couple of minutes.

### 3.2 BRD application example

The web tool based on the BRD model is produced from the YAML specification as explained above (2.4). The version presented here is provided together with the PASTE software as an illustration of what can be done with this tool generator, and the resulting web application has been presented and discussed with researchers in the veterinarian field. The tool is already fully functional, and we already planned to present it in focus groups with farmers and field veterinarians to identify feature improvements.

Currently, the BRD tool starts with a first form to specify on-farm situation (Fig. 5), built from the YAML specification (Fig. 6). This form accounts for farming practices such as batch composition (in terms of risk levels, batch allocation), population (batch size and number of batches), dominant pathogens, and economic costs.

After choosing scenarios to compare (here, using risk level or not to allocate beef calves to batches — see Fig. 8-9), the user can indicate execution settings (such as the duration of the simulation — see SI Fig. 1) and launch the simulations. After the stochastic repetitions of all scenarios are finished and their outputs processed, statistics are provided to the user, for instance a comparison between the distribution of antimicrobial doses at batch scale depending on how young beef bulls have been allocated to batches (Fig. 10, built from the specification of Fig.11), or the average dynamics of severe BRD cases at farm scale in both scenarios (Fig. SI 3, built from Fig. SI-4).

## 4. Discussion

### 4.1 Flexibility and genericity

To promote the use of decision support tools based on mechanistic models in epidemiology, we advocate for automating the transformation chain to produce web DST from existing mechanistic epidemiological models. The EMULSION-PASTE software pipeline is a first operational implementation of this approach. It fosters co-design between modellers and other scientists, facilitates model assessment and revisions, reduces the risk of errors in the simulation code (shared within the simulation engine).

A methodological principle underlies the whole EMULSION-PASTE approach: the separation of concerns (Picault et al., 2017). Modellers are in charge of designing and assessing the mechanistic models, while tool designers and stakeholders specify their needs. This ensures that the persons involved in each task have the proper skills. Hence, the model can be as complex as necessary to address multiple research questions, while tools can provide more focused views and incorporate additional concerns (e.g., cost/benefit considerations, realistic parameterization to account for field situations). This principle is implemented through a separation between the concepts (model/tool structure and components, role, and values of parameters, etc.) and the code (simulation architecture and algorithms, web application). This is made possible using methods at the crossroads between AI and software engineering: knowledge representation (DSL) and generic software patterns (multi-level agent-based architecture, web framework such as Django). Besides, this separation between model description and the simulation code is crucial to make web code generation possible on top of existing mechanistic models.

The main added value of this approach is that revisions of the tool can be made in a very reactive way when new features are required, as no additional development effort is needed. The changes must be incorporated in the tool specification, then regenerating the code is immediate. Also, the web application benefits from model updates, such as new knowledge on the modelled system, an increased accuracy of parameter values, or model components such as new control measures.

The development of the web application does not require to write any code, which also reduces the time spent in chasing bugs. The reliability of the application is based upon the generic software chain, which is used (hence, tested and corrected) in many diverse situations to generate a diversity of web tools. The open-source nature of EMULSION and PASTE also accelerates the identification and correction of potential bugs by the community of software users. Also, customizing the tools produced by PASTE to go from the beta version to a final product can be done by any web developer, as the web architecture relies on broadly used technologies (i.e. Django and R-Shiny).

### 4.2 Alternative approaches

So far and to the best of our knowledge, there is no equivalent approach to foster such continuum between academic research in animal health and on-farm applications. Regarding epidemiological modelling, DSLs have been proposed in only two platforms to represent mechanistic models: Kendrick (Bui T. et al., 2019) and EMULSION (Picault et al., 2019). In Kendrick, the DSL is a simplified version of the Smalltalk programming language and has a procedural syntax, i.e. the modellers have to provide instructions in the proper order to build the model. As a result, it is still very close to a classical high-level programming language. In contrast, in EMULSION the DSL is declarative, i.e. model components can be provided in any order as long as they organize in a consistent manner. SimInf (Widgren et al., 2019) is a R library that makes it possible to integrate with existing epidemiological data and animal movements, but the lack of associated DSL hinders the capability to design complex epidemiological models. The Java framework BROADWICK (O’Hare et al., 2016) can tackle complex epidemiological models at large scale but requires programming. Other simulation software can bring intrinsic limitations to the range of users, as they often require technical skills for installation, configuration, and execution of modelling programs, which are rarely mastered by decision makers. Wrapping a modelling software into a graphical user interface, such as a stand-alone graphical program or a web front-end, reduces the technicity of software usage but does not address the issue of the complexity of the model itself. An alternative consists in keeping model complexity low (by providing a small set of predefined model structures) to foster the exploration of scenarios, relying upon integration with external data and geographical information systems to enhance the realism of the simulations (Broeck et al., 2011). However, this approach assumes that the users of the web tool have enough modelling skills to understand the limitations of the underlying models and avoid overinterpretations of the results. Similarly, some general-purpose simulation platforms such as NetLogo (Wilensky and Rand, 2015), Repast (North et al., 2013) or GAMA (Grignard et al., 2013), provide great user interfaces, the capability to connect to geographical information systems, and a high-level programming language. However, installing and configuring those platforms, developing epidemiological models usable in real situations, and running simulations with relevant outputs again requires technical skills in modelling and computer science which makes them difficult to be used by health managers in an autonomous way.

As academic models are not directly usable as DST by non-modeller end-users, participatory methods (Voinov et al., 2018; Voinov and Bousquet, 2010) aim at the co-design of a model and the associated DST at the same time, by modellers and stakeholders, for a given practical question, often resulting in an improved engagement by stakeholders in the process and higher confidence in model outcomes (Lee et al., 2022). This however results in a model that focuses on a narrow application-oriented question, e.g. assessing the cost-benefit trade-off of control measures (Bennett et al., 2010), and thus limits the scope of research questions that can be addressed, though the development time is almost as long as for a broader model.

### 4.3 Limitations

There are currently two main limitations in our approach. The first one is that performance issues may arise, depending on the model complexity, but also on the technology used for the deployment (computing power of the calculation server, limited performance of EMULSION which is also written in Python).

The second one is that, so far, the PASTE approach is based on models written with EMULSION. However, the software could be adapted to any other simulation software. To do so, the tool developer should write a YAML document mimicking the sections of an EMULSION model that describe the list and default values of parameters, and outputs available produced by the simulation (assuming they are made available as a CSV file as in EMULSION). Then, the developer should specify the syntax for running the simulation program (instead of EMULSION) with proper parameter values. This can be done in the YAML file in a section called “emulsion_syntax” (L20-25 in SI section 2). Another way to proceed is to use a script to mimic the EMULSION command syntax and transform options into the original syntax of the target simulation program.

## 5 Conclusions

The EMULSION-PASTE coupling involved symbolic AI methods, such as knowledge representation and agent-based simulation architectures, including multi-level simulation and organizational systems. AI is combined with software engineering, especially model-driven methods to define a meta-model and domain-specific languages to formalize the description of the system to tackle.

AI does not make the complexity of designing epidemiological models and DST “vanish”: it rather helps splitting the complexity in several sub-domains through a separation of concerns: designing the conceptual model, designing a relevant tool for end-users, implementing the model within EMULSION DSL, implementing the tool specifications within PASTE DSL, and of course incorporate in EMULSION and PASTE the algorithms and data structures which are required to do so. Each step is more focused on a single set of skills.

This work is also aimed at facilitating the appropriation of academic research by stakeholders through an acceleration of the development of flexible web tools based on academic models. To go further, we are currently integrating the PASTE software into a model-driven engineering software, JetBrains MPS (Meta-Programming System), to provide a dedicated standalone editor, featuring user-ready templates, completion, consistency verification, and more flexible code generation which should result in an enhanced experience for the tool developers.

In the longer term, we plan to extend PASTE language and engine in two directions. First, we consider making it possible to connect web tools with data, which could for instance be used to update initial conditions, either from prior observation data, or from real-time information flow (Tedeschi et al., 2021). That would indeed enhance the realism of situations that the tools can handle and relieve users from updating the information themselves. Second, as scenarios are based on the combination of several components, the system could automatically identify the most efficient scenarios in respect of a given metric, and propose an explicit intervention ranking, which would provide not only a factual and “neutral” comparison between scenarios as in PASTE so far, but explicit and intelligible recommendations.

## Supporting information

paste.yaml

## Availability of data and materials

EMULSION is an open-source software distributed as a Python module (https://sourcesup.renater.fr/www/emulsion-public). PASTE is also provided as an open-source software, available from the following public GIT repository: https://forgemia.inrae.fr/dynamo/software/PASTE-public-release.

## Funding

This work was carried out with the financial support of INRAE, Partnerships and Innovation Transfer Division (ATOM project) and with a grant of the Carnot institute France Futur Élevage (project SEPTIME). This work has received funding from the European Union’s Horizon 2020 research and innovation programme under grant agreement No. 101000494 (DECIDE). This publication only reflects the authors’ views and the Research Executive Agency is not responsible for any use that may be made of the information it contains. This work was also supported by the French region Pays de la Loire (PULSAR grant). No funder was involved in the scientific or editorial choices made for this study.

## Acknowledgements

We thank our colleagues from the BIOEPAR Dynamo team, and Dr. Massimo Tisi (IMT Atlantique, Nantes, France), for their comments and advice on this work.

## Author’s contributions

Conception and design of the study: SP, PE, VS, SA. Creation of software model: GN, SP, VS. Design of example tools: SP, SA, GN, PE, BS. Drafting of the manuscript: SP. Revisions of the manuscript: ALL. All authors read and approved the final manuscript.

## Competing interests

The authors declare that they have no competing interests.

## Footnotes

1 This article is currently under moderate revision, for publication in PVM.

## Notes

### Competing Interest Statement

The authors have declared no competing interest.

## References

Amouroux, E., Desvaux, S., Drogoul, A., 2008. Towards Virtual Epidemiology: An Agent-Based Approach to the Modeling of H5N1 Propagation and Persistence in North-Vietnam, in: Bui, T.D., Ho, T.V., Ha, Q.T. (Eds.), 11th Pacific Rim Int. Conf. on Multi-Agents (PRIMA), Lecture Notes in Computer Science. Springer, pp. 26–33. 10.1007/978-3-540-89674-6_6

Barton, C.M., Alberti, M., Ames, D., Atkinson, J.-A., Bales, J., Burke, E., Chen, M., Diallo, S.Y., Earn, D.J.D., Fath, B., Feng, Z., Gibbons, C., Hammond, R., Heffernan, J., Houser, H., Hovmand, P.S., Kopainsky, B., Mabry, P.L., Mair, C., Meier, P., Niles, R., Nosek, B., Osgood, N., Pierce, S., Polhill, J.G., Prosser, L., Robinson, E., Rosenzweig, C., Sankaran, S., Stange, K., Tucker, G., 2020. Call for transparency of COVID-19 models. Science 368, 482.2-483. 10.1126/science.abb8637

Beaunée, G., Vergu, E., Joly, A., Ezanno, P., 2017. Controlling bovine paratuberculosis at a regional scale: Towards a decision modelling tool. J. Theor. Biol. 435, 157–183. 10.1016/j.jtbi.2017.09.012

Ben-Kiki, O., Evans, C., döt Net, I., 2009. YAML Ain’t Markup Language (YAML™) version 1.2.

Bennett, R., McClement, I., McFarlane, I., 2010. An economic decision support tool for simulating paratuberculosis control strategies in a UK suckler beef herd. Prev. Vet. Med. 93, 286–293. 10.1016/j.prevetmed.2009.11.006

Bigley Dunham, J., 2005. An Agent-Based Spatially Explicit Epidemiological Model in MASON. J. Artif. Soc. Soc. Simul. 9.

Brock, J., Lange, M., More, S.J., Graham, D., Thulke, H.-H., 2020. Reviewing age-structured epidemiological models of cattle diseases tailored to support management decisions: Guidance for the future. Prev. Vet. Med. 174, 104814. 10.1016/j.prevetmed.2019.104814

Broeck, W.V. den, Gioannini, C., Gonçalves, B., Quaggiotto, M., Colizza, V., Vespignani, A., 2011. The GLEaMviz computational tool, a publicly available software to explore realistic epidemic spreading scenarios at the global scale. BMC Infect. Dis. 11. 10.1186/1471-2334-11-37

Bucchiarone, A., Ciccozzi, F., Lambers, L., Pierantonio, A., Tichy, M., Tisi, M., Wortmann, A., Zaytsev, V., 2021. What Is the Future of Modeling? IEEE Softw. 38, 119–127. 10.1109/MS.2020.3041522

Bui T. M.A., Papoulias, N., Stinckwich, S., Ziane, M., Roche, B., 2019. The Kendrick modelling platform: language abstractions and tools for epidemiology. BMC Bioinformatics 20, 312. 10.1186/s12859-019-2843-0

Danon, L., Brooks-Pollock, E., Bailey, M., Keeling, M., 2021. A spatial model of COVID-19 transmission in England and Wales: early spread, peak timing and the impact of seasonality. Philos. Trans. R. Soc. B Biol. Sci. 376, 20200272. 10.1098/rstb.2020.0272

Del Valle, S.Y., McMahon, B.H., Asher, J., Hatchett, R., Lega, J.C., Brown, H.E., Leany, M.E., Pantazis, Y., Roberts, D.J., Moore, S., Peterson, A.T., Escobar, L.E., Qiao, H., Hengartner, N.W., Mukundan, H., 2018. Summary results of the 2014-2015 DARPA Chikungunya challenge. BMC Infect. Dis. 18, 245. 10.1186/s12879-018-3124-7

Divya, K.L., Mhatre, P.H., Venkatasalam, E.P., Sudha, R., 2021. Crop Simulation Models as Decision-Supporting Tools for Sustainable Potato Production: a Review. Potato Res. 64, 387–419. 10.1007/s11540-020-09483-9

Dykes, J., Abdul-Rahman, A., Archambault, D., Bach, B., Borgo, R., Chen, M., Enright, J., Fang, H., Firat, E.E., Freeman, E., Gönen, T., Harris, C., Jianu, R., John, N.W., Khan, S., Lahiff, A., Laramee, R.S., Matthews, L., Mohr, S., Nguyen, P.H., Rahat, A.A.M., Reeve, R., Ritsos, P.D., Roberts, J.C., Slingsby, A., Swallow, B., Torsney-Weir, T., Turkay, C., Turner, R., Vidal, F.P., Wang, Q., Wood, J., Xu, K., 2022. Visualization for epidemiological modelling: challenges, solutions, reflections and recommendations. Philos. Trans. R. Soc. Math. Phys. Eng. Sci. 380, 20210299. 10.1098/rsta.2021.0299

Ezanno, P., Andraud, M., Beaunée, G., Hoch, T., Krebs, S., Rault, A., Touzeau, S., Vergu, E., Widgren, S., 2020. How mechanistic modelling supports decision making for the control of enzootic infectious diseases. Epidemics 100398. 10.1016/j.epidem.2020.100398

Ezanno, P., Picault, S., Bareille, S., Beaunée, B., Jan Boender, G., Dankwa, E.A., Deslandes, F., Donnelly, C.A., Hagenaars, T.J., Hayes, S., Jori, F., Lambert, S., Mancini, M., Munoz, F., Pleydell, D.R.J., Thompson, R.N., Vergu, E., Vignes, M., Vergne, T., 2022. The African swine fever modelling challenge: model comparison and lessons learnt. Epidemics 100615. 10.1016/j.epidem.2022.100615

Ezanno, P., Picault, S., Beaunée, G., Bailly, X., Muñoz, F., Duboz, R., Monod, H., Guégan, J.-F., 2021. Research perspectives on animal health in the era of artificial intelligence. Vet. Res. 52, 40. 10.1186/s13567-021-00902-4

Ferguson, N.M., Cummings, D.A.T., Cauchemez, S., Fraser, C., Riley, S., Meeyai, A., Iamsirithaworn, S., Burke, D.S., 2005. Strategies for containing an emerging influenza pandemic in Southeast Asia. Nature 437, 209–214. 10.1038/nature04017

Grignard, A., Taillandier, P., Gaudou, B., Vo, D.A., Huynh, N.Q., Drogoul, A., 2013. GAMA 1.6: Advancing the Art of Complex Agent-Based Modeling and Simulation, in: Hutchison, D., Kanade, T., Kittler, J., Kleinberg, J.M., Mattern, F., Boella, G., Elkind, E., Savarimuthu, B.T.R., Dignum, F., Purvis, M.K. (Eds.), PRIMA 2013: Principles and Practice of Multi-Agent Systems. Springer Berlin Heidelberg, Berlin, Heidelberg, pp. 117–131. 10.1007/978-3-642-44927-7_9

Halloran, M.E., Ferguson, N.M., Eubank, S., Longini, I.M., Cummings, D.A.T., Lewis, B., Xu, S., Fraser, C., Vullikanti, A., Germann, T.C., Wagener, D., Beckman, R., Kadau, K., Barrett, C., Macken, C.A., Burke, D.S., Cooley, P., 2008. Modeling targeted layered containment of an influenza pandemic in the United States. Proc. Natl. Acad. Sci. 105, 4639–4644. 10.1073/pnas.0706849105

Hethcote, H.W., 2000. The Mathematics of Infectious Diseases. SIAM Rev. 42, 599–653. 10.1137/S0036144500371907

Hinch, R., Probert, W.J.M., Nurtay, A., Kendall, M., Wymant, C., Hall, M., Lythgoe, K., Bulas Cruz, A., Zhao, L., Stewart, A., Ferretti, L., Montero, D., Warren, J., Mather, N., Abueg, M., Wu, N., Legat, O., Bentley, K., Mead, T., Van-Vuuren, K., Feldner-Busztin, D., Ristori, T., Finkelstein, A., Bonsall, D.G., Abeler-Dörner, L., Fraser, C., 2021. OpenABM-Covid19— An agent-based model for non-pharmaceutical interventions against COVID-19 including contact tracing. PLOS Comput. Biol. 17, e1009146. 10.1371/journal.pcbi.1009146

Johansson, M.A., Apfeldorf, K.M., Dobson, S., Devita, J., Buczak, A.L., Baugher, B., Moniz, L.J., Bagley, T., Babin, S.M., Guven, E., Yamana, T.K., Shaman, J., Moschou, T., Lothian, N., Lane, A., Osborne, G., Jiang, G., Brooks, L.C., Farrow, D.C., Hyun, S., Tibshirani, R.J., Rosenfeld, R., Lessler, J., Reich, N.G., Cummings, D.A.T., Lauer, S.A., Moore, S.M., Clapham, H.E., Lowe, R., Bailey, T.C., García-Díez, M., Carvalho, M.S., Rodó, X., Sardar, T., Paul, R., Ray, E.L., Sakrejda, K., Brown, A.C., Meng, X., Osoba, O., Vardavas, R., Manheim, D., Moore, M., Rao, D.M., Porco, T.C., Ackley, S., Liu, F., Worden, L., Convertino, M., Liu, Y., Reddy, A., Ortiz, E., Rivero, J., Brito, H., Juarrero, A., Johnson, L.R., Gramacy, R.B., Cohen, J.M., Mordecai, E.A., Murdock, C.C., Rohr, J.R., Ryan, S.J., Stewart-Ibarra, A.M., Weikel, D.P., Jutla, A., Khan, R., Poultney, M., Colwell, R.R., Rivera-García, B., Barker, C.M., Bell, J.E., Biggerstaff, M., Swerdlow, D., Mier-y-Teran-Romero, L., Forshey, B.M., Trtanj, J., Asher, J., Clay, M., Margolis, H.S., Hebbeler, A.M., George, D., Chretien, J.-P., 2019. An open challenge to advance probabilistic forecasting for dengue epidemics. Proc. Natl. Acad. Sci. 201909865. 10.1073/pnas.1909865116

Keeling, M., 2005. The implications of network structure for epidemic dynamics. Theor. Popul. Biol. 67, 1–8. 10.1016/j.tpb.2004.08.002

Keeling, M.J., Rohani, P., 2008. Modeling Infectious Diseases in Humans and Animals. Princeton University Press.

Kerr, C.C., Stuart, R.M., Mistry, D., Abeysuriya, R.G., Rosenfeld, K., Hart, G.R., Núñez, R.C., Cohen, J.A., Selvaraj, P., Hagedorn, B., George, L., Jastrzębski, M., Izzo, A.S., Fowler, G., Palmer, A., Delport, D., Scott, N., Kelly, S.L., Bennette, C.S., Wagner, B.G., Chang, S.T., Oron, A.P., Wenger, E.A., Panovska-Griffiths, J., Famulare, M., Klein, D.J., 2021. Covasim: An agent-based model of COVID-19 dynamics and interventions. PLOS Comput. Biol. 17, e1009149. 10.1371/journal.pcbi.1009149

Lee, G.Y., Hickie, I.B., Occhipinti, J.-A., Song, Y.J.C., Skinner, A., Camacho, S., Lawson, K., Hilber, A.M., Freebairn, L., 2022. Presenting a comprehensive multi-scale evaluation framework for participatory modelling programs: A scoping review. PLOS ONE 17, e0266125. 10.1371/journal.pone.0266125

Mathieu, P., Morvan, G., Picault, S., 2018. Multi-level agent-based simulations: Four design patterns. Simul. Model. Pract. Theory 83, 51–64. 10.1016/j.simpat.2017.12.015

Meredith, H.R., Arehart, E., Grantz, K.H., Beams, A., Sheets, T., Nelson, R., Zhang, Y., Vinik, R.G., Barfuss, D., Pettit, J.C., McCaffrey, K., Dunn, A.C., Good, M., Frattaroli, S., Samore, M.H., Lessler, J., Lee, E.C., Keegan, L.T., 2021. Coordinated Strategy for a Model-Based Decision Support Tool for Coronavirus Disease, Utah, USA. Emerg. Infect. Dis. 27, 1259–1265. 10.3201/eid2705.203075

Muellner, U., Fournié, G., Muellner, P., Ahlstrom, C., Pfeiffer, D.U., 2018. epidemix —An interactive multi-model application for teaching and visualizing infectious disease transmission. Epidemics 23, 49–54. 10.1016/j.epidem.2017.12.003

North, M.J., Collier, N.T., Ozik, J., Tatara, E.R., Macal, C.M., Bragen, M., Sydelko, P., 2013. Complex adaptive systems modeling with Repast Simphony. Complex Adapt. Syst. Model. 1, 3. 10.1186/2194-3206-1-3

O’Hare, A., Lycett, S.J., Doherty, T., M. Salvador, L.C., Kao, R.R., 2016. Broadwick: a framework for computational epidemiology. BMC Bioinformatics 17. 10.1186/s12859-016-0903-2

Peng, R.D., 2011. Reproducible Research in Computational Science. Science 334, 1226–1227. 10.1126/science.1213847

Picault, S., Huang, Y.-L., Sicard, V., Arnoux, S., Beaunée, G., Ezanno, P., 2019. EMULSION: Transparent and flexible multiscale stochastic models in human, animal and plant epidemiology. PLOS Comput. Biol. 15, e1007342. 10.1371/journal.pcbi.1007342

Picault, S., Huang, Y.-L., Sicard, V., Ezanno, P., 2017. Enhancing Sustainability of Complex Epidemiological Models through a Generic Multilevel Agent-based Approach, in: Sierra, C. (Ed.), Proceedings of the 26th International Joint Conference on Artificial Intelligence (IJCAI’2017). AAAI, Melbourne, Australia. 10.24963/ijcai.2017/53

Reguly, I.Z., Csercsik, D., Juhász, J., Tornai, K., Bujtár, Z., Horváth, G., Keömley-Horváth, B., Kós, T., Cserey, G., Iván, K., Pongor, S., Szederkényi, G., Röst, G., Csikász-Nagy, A., 2022. Microsimulation based quantitative analysis of COVID-19 management strategies. PLOS Comput. Biol. 18, e1009693. 10.1371/journal.pcbi.1009693

Reich, N.G., Brooks, L.C., Fox, S.J., Kandula, S., McGowan, C.J., Moore, E., Osthus, D., Ray, E.L., Tushar, A., Yamana, T.K., Biggerstaff, M., Johansson, M.A., Rosenfeld, R., Shaman, J., 2019. A collaborative multiyear, multimodel assessment of seasonal influenza forecasting in the United States. Proc. Natl. Acad. Sci. 116, 3146–3154. 10.1073/pnas.1812594116

Roche, B., Guegan, J.-F., Bousquet, F., 2008. Multi-agent systems in epidemiology: a first step for computational biology in the study of vector-borne disease transmission. BMC Bioinformatics 9, 435. 10.1186/1471-2105-9-435

Russell, S.J., Norvig, P., 2010. Artificial Intelligence. A Modern Approach, 3rd edition. ed. Upper Saddle River, New Jersey.

Schneider, K.A., Ngwa, G.A., Schwehm, M., Eichner, L., Eichner, M., 2020. The COVID-19 pandemic preparedness simulation tool: CovidSIM. BMC Infect. Dis. 20, 859. 10.1186/s12879-020-05566-7

Shea, K., Borchering, R.K., Probert, W.J.M., Howerton, E., Bogich, T.L., Li, S.-L., Van Panhuis, W.G., Viboud, C., Aguás, R., Belov, A.A., Bhargava, S.H., Cavany, S.M., Chang, J.C., Chen, C., Chen, J., Chen, S., Chen, Y., Childs, L.M., Chow, C.C., Crooker, I., Del Valle, S.Y., España, G., Fairchild, G., Gerkin, R.C., Germann, T.C., Gu, Q., Guan, X., Guo, L., Hart, G.R., Hladish, T.J., Hupert, N., Janies, D., Kerr, C.C., Klein, D.J., Klein, E.Y., Lin, G., Manore, C., Meyers, L.A., Mittler, J.E., Mu, K., Núñez, R.C., Oidtman, R.J., Pasco, R., Pastore Y Piontti, A., Paul, R., Pearson, C.A.B., Perdomo, D.R., Perkins, T.A., Pierce, K., Pillai, A.N., Rael, R.C., Rosenfeld, K., Ross, C.W., Spencer, J.A., Stoltzfus, A.B., Toh, K.B., Vattikuti, S., Vespignani, A., Wang, L., White, L.J., Xu, P., Yang, Y., Yogurtcu, O.N., Zhang, W., Zhao, Y., Zou, D., Ferrari, M.J., Pannell, D., Tildesley, M.J., Seifarth, J., Johnson, E., Biggerstaff, M., Johansson, M.A., Slayton, R.B., Levander, J.D., Stazer, J., Kerr, J., Runge, M.C., 2023. Multiple models for outbreak decision support in the face of uncertainty. Proc. Natl. Acad. Sci. 120, e2207537120. 10.1073/pnas.2207537120

Sicard, V., Andraud, M., Picault, S., 2022a. Coupling spatial and temporal structure in batch rearing modeling for understanding the spread of the swine influenza A virus, in: Proceedings of the Conference of the Society for Veterinary Epidemiology and Preventive Medicine (SVEPM). Belfast, UK.

Sicard, V., Andraud, M., Picault, S., 2022b. A declarative modelling language for the design of complex structured agent-based epidemiological models, in: Proceedings of the 20th International Conference on Practical Applications of Agents and Multi-Agent Systems (PAAMS), Lecture Notes in Computer Science. Presented at the PAAMS 2022, Springer, L’Aquila, Italia, pp. 385–396. 10.1007/978-3-031-18192-4_31

Sicard, V., Andraud, M., Picault, S., 2021. Organization as a multi-level design pattern for agent-based simulation of complex systems, in: Proceedings of the 13th International Conference on Agents and Artificial Intelligence (ICAART’2021). SCITEPRESS, pp. 232–241. 10.5220/0010223202320241

Sorin-Dupont, B., Picault, S., Pardon, B., Ezanno, P., Assié, S., 2023. Modeling the effects of farming practices on bovine respiratory disease in a multi-batch cattle fattening farm. Prev. Vet. Med.

Sutherland, W.J., Freckleton, R.P., 2012. Making predictive ecology more relevant to policy makers and practitioners. Philos. Trans. R. Soc. B Biol. Sci. 367, 322–330. 10.1098/rstb.2011.0181

Tedeschi, L.O., Greenwood, P.L., Halachmi, I., 2021. Advancements in sensor technology and decision support intelligent tools to assist smart livestock farming. J. Anim. Sci. 99, skab038. 10.1093/jas/skab038

Thulke, H.-H., 2011. Application of recent approaches in modelling for Animal Health. Prev. Vet. Med. 99, 1–3. 10.1016/j.prevetmed.2011.01.007

Thulke, H.-H., Lange, M., Tratalos, J.A., Clegg, T.A., McGrath, G., O’Grady, L., O’Sullivan, P., Doherty, M.L., Graham, D.A., More, S.J., 2018. Eradicating BVD, reviewing Irish programme data and model predictions to support prospective decision making. Prev. Vet. Med. 150, 151–161. 10.1016/j.prevetmed.2017.11.017

Viboud, C., Sun, K., Gaffey, R., Ajelli, M., Fumanelli, L., Merler, S., Zhang, Q., Chowell, G., Simonsen, L., Vespignani, A., 2018. The RAPIDD ebola forecasting challenge: Synthesis and lessons learnt. Epidemics 22, 13–21. 10.1016/j.epidem.2017.08.002

Voinov, A., Bousquet, F., 2010. Modelling with stakeholders. Environ. Model. Softw. 25, 1268–1281. 10.1016/j.envsoft.2010.03.007

Voinov, A., Jenni, K., Gray, S., Kolagani, N., Glynn, P.D., Bommel, P., Prell, C., Zellner, M., Paolisso, M., Jordan, R., Sterling, E., Schmitt Olabisi, L., Giabbanelli, P.J., Sun, Z., Le Page, C., Elsawah, S., BenDor, T.K., Hubacek, K., Laursen, B.K., Jetter, A., Basco-Carrera, L., Singer, A., Young, L., Brunacini, J., Smajgl, A., 2018. Tools and methods in participatory modeling: Selecting the right tool for the job. Environ. Model. Softw. 109, 232–255. 10.1016/j.envsoft.2018.08.028

Webb, C.T., Ferrari, M., Lindström, T., Carpenter, T., Dürr, S., Garner, G., Jewell, C., Stevenson, M., Ward, M.P., Werkman, M., Backer, J., Tildesley, M., 2017. Ensemble modelling and structured decision-making to support Emergency Disease Management. Prev. Vet. Med. 138, 124–133. 10.1016/j.prevetmed.2017.01.003

Widgren, S., Bauer, P., Eriksson, R., Engblom, S., 2019. SimInf: An R Package for Data-Driven Stochastic Disease Spread Simulations. J. Stat. Softw. 91. 10.18637/jss.v091.i12

Wilensky, U., Rand, W., 2015. An introduction to agent-based modeling: modeling natural, social, and engineered complex systems with NetLogo. The MIT Press, Cambridge, Massachusetts.

