## Supporting Information for "Leveraging artificial intelligence and software engineering methods in epidemiology for the co-creation of decision-support tools based on mechanistic models"

**Sébastien Picault*[1], Guita Niang[1], Vianney Sicard[1], Baptiste Sorin-Dupont[1], Sébastien Assié[1], Pauline Ezanno[1]**

1 A full example of an PASTE-generated web application ........ 1
2 PASTE tool specification (YAML file) for the BRD example ........ 7

### 1 A full example of an PASTE-generated web application

This section presents additional screenshots from the web tool on Bovine Respiratory Disease (main manuscript sections 2.5 and 3.2) with the underlying YAML specification.

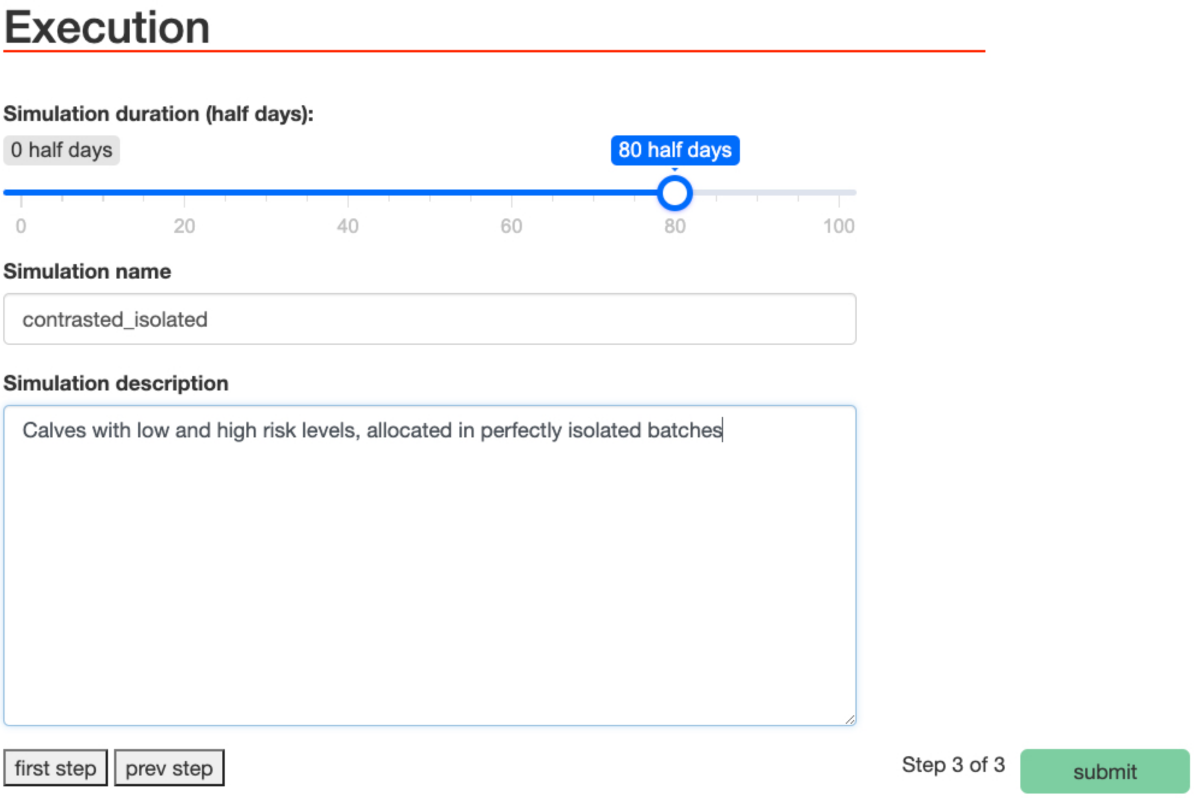


*Figure SI1. Input form for simulation execution. The user can indicate the duration of the period to simulate, associated with a simulation name and comments. The form specifications are defined in the YAML file L80-192 (Fig. SI2).*

```
input_forms:
...
   - execution:
       - total_duration:
           type: IntegerField
           label: "Simulation duration (half days)"
           required: True
           widget: NumberInput
           widget_attrs:
             'data-min': 0
             'data-step': 2
             'data-value': 60
             'data-skin': 'round'
             'data-label': 'half days'
             'data-grid': 'true'
             'data-grid-num': 5
             'data-max': 100
       - simulation_name:
           type: CharField
           label: "Simulation name"
           required: True
           widget: forms.TextInput

       - simulation_description:
           type: CharField
           label: "Simulation description"
           required: False
           widget: forms.Textarea
```

*Figure SI2. Specification of the execution input form (Fig. SI1)*

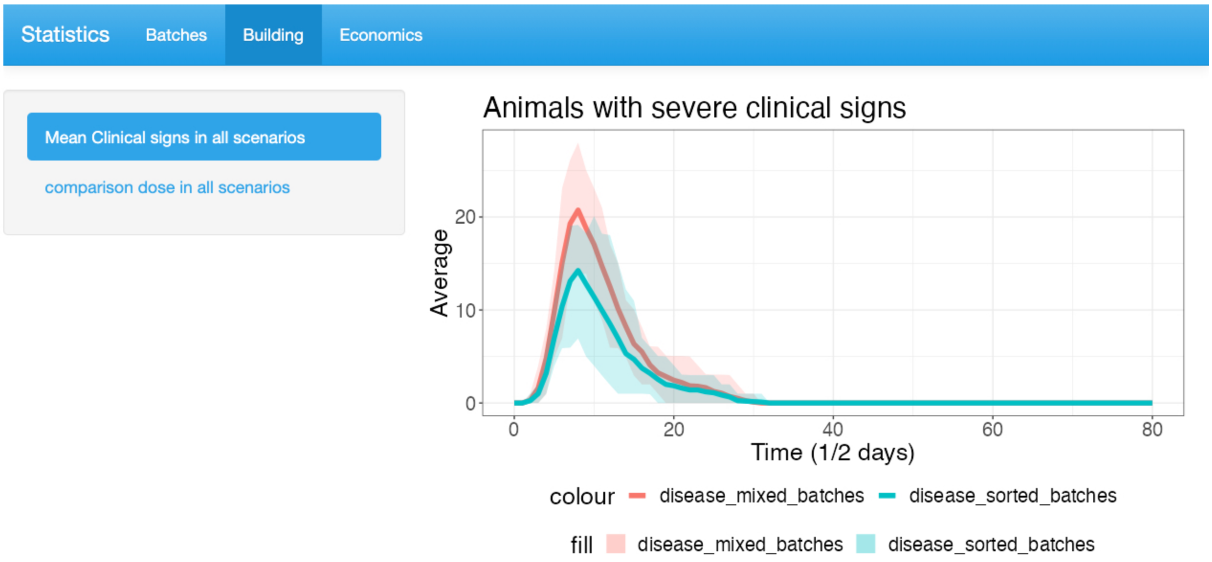


*Figure SI3.Output page showing results at farm scale over time. The figure presents the number of severely diseased animals (average and 5th-95th percentiles) over time in two scenarios. The graph definition is L255-280, based on the treatment of simulation results specified in L195-207 (Fig. SI4).*

```
statistics:
  for_each_scenario:
    - mean_C:
        - desc: "Animals with severe clinical signs (mean, q05, q95)'
        - target_variable: [ "metapop_total_C" ]
        - groupby: [ "step" ]
        - summarise:
            - mean
            - perc_0.5:
                value: "quantile(..., 0.05)"
                desc: "5th percentile"
            - perc_0.95:
                value: "quantile(..., 0.95)"
                desc: "95th percentile"
.......................................................................
graphics:
  plot_mean_C:
    - data: "mean_C"
    - on_page: ["Building"]
    - label: ["Mean Clinical signs in all scenarios"]
    - single_graph: "yes"
    - scenario: ["disease_mixed_batches", "disease_sorted_batches"]
    - plot_variables:
        x: "step"
        y: "mean_metapop_total_C"
    - plot_type:
        - line:
            plot_variables:
              colour: "scenario_name"
            plot_options:
              linewidth: "2"
        - ribbon:
            plot_variables:
              ymax: "perc_0.95_metapop_total_C"
              ymin: "perc_0.5_metapop_total_C"
              fill: "scenario_name"
            plot_options:
              alpha: "0.2"
    - plot_annotations:
        title: "Animals with severe clinical signs"
        x_title: "Time (1/2 days)"
        y_title: "Average"
```

*Figure SI4. Specification of the plot (Fig. SI3)*

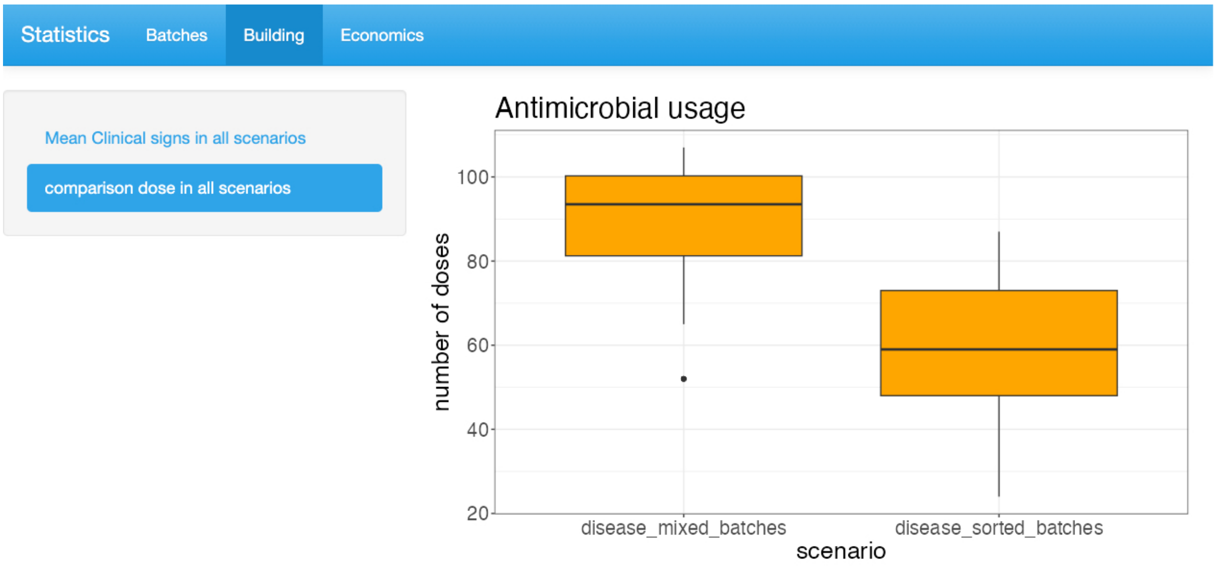


*Figure SI5. Output page showing AMU at farm scale. The figure presents the number of antimicrobial doses used by all batches (distribution over the stochastic repetitions) in two scenarios (left vs. right). The graph definition is L358-371, based on the treatment of simulation results specified in L208-211 (Fig. SI6).*

```
194 statistics:
...........................................................................
208     - nb_doses:
209         - desc: "Number of doses at farm scale"
210         - target_variable: [ "metapop_nb_doses" ]
211         - condition: [ 'step == MAX.STEP' ]
...........................................................................
254 graphics:
...........................................................................
358   plot_doses:
359     - data: "nb_doses"
360     - on_page: ["Building"]
361     - label: ["comparison dose in all scenarios"]
362     - single_graph: "yes"
363     - scenario: ["disease_mixed_batches", "disease_sorted_batches"]
364     - plot_variables:
365         y: "metapop_nb_doses"
366     - plot_type:
367         - boxplot:
368             plot_variables:
369               x: "scenario_name"
370             plot_options:
371               fill: "orange"
```

*Figure SI6. Specification of the plot (Fig. SI5).*

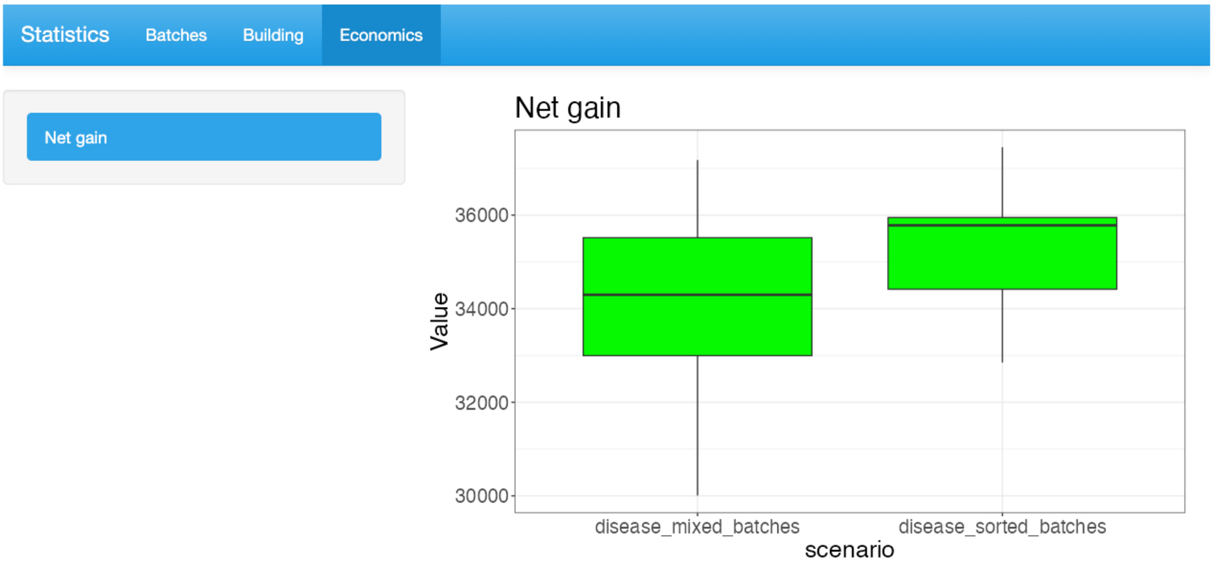


*Figure SI7. Output page showing economic gain at farm scale. The figure presents the net gain of fattening at farm scale (distribution over the stochastic repetitions) in two scenarios (left vs. right). The graph definition is L388-390, based on the treatment of simulation results specified in L239-246 (Fig. SI8).*

```
194 statistics:
.......................................................................
239     - economics:
240         - desc: "Net gain of fattening"
241         - target_variable: ["total_population", "metapop_nb_doses", '
242         - condition: [ 'step == MAX.STEP' ]
243         - add_new_column:
244             gain:
245               value: "selling_price_bull *  total_population - cost_
246               desc: "Net gain"
.......................................................................
254 graphics:
.......................................................................
378   plot_gains:
379     - data: "economics"
380     - on_page: ["Economics"]
381     - label: ["Net gain"]
382     - single_graph: "yes"
383     - scenario: ["disease_mixed_batches", "disease_sorted_batches"]
384     - plot_variables:
385         y: "gain"
386     - plot_type:
387         - boxplot:
388             plot_variables:
389               x: "scenario_name"
390             plot_options:
```

*Figure SI8 Specification of the plot (Fig. SI7).*

### 2 PASTE tool specification (YAML file) for the BRD example

This file is available on the public PASTE GIT repository (https://forgemia.inrae.fr/dynamo/software/PASTE-public-release), in directory "PASTE_examples/brd_example".

```yaml
django_project_name: brd_project

django_app_name: brd_app

input_files_path:
  path: brd_tool_specification/input_files

model_emulsion_file:
  file_name: brd-svepm-paste.yaml

emulsion_output_directory:
  path: outputs/

emulsion_setup:
  model_name: BRD
  number_of_repetitions: 20
  figure_text_size: 20

  emulsion_syntax:
    command: 'emulsion run'
    runs: '-r'
    outputs: '--output-dir'
    options: '--silent'
    params: '-p'

show_hidden_parameters: no

calcul_URI: "http://127.0.0.1:8100/"

input_parameters:
  - block_values:
      pathogen:
        - file_path: "input_files/pathogens.csv"
      batch_isolation:
        - file_path: "input_files/biosecurity.csv"
      risk_level:
        - file_path: "input_files/risk_level.csv"

  - user_parameters_values:
      - cost_of_antiobiotic_dose: 10
      - cost_purchase_calf: 950
      - selling_price_bull: 1330
      - total_feeding_cost: 312
      - daily_disease_impact: 0.2
      - batch_size: 10
      - number_batches: 3

  - user_parameters_matrices: []

scenarios_components:
  - health_conditions:
      - no_disease:
          optional: "no"
```

```yaml
          proportion_carrier_low: 0
          proportion_carrier_medium: 0
          proportion_carrier_high: 0
      - disease:
          optional: "yes"

  - batch_composition:
      - mixed_batches:
          optional: "yes"
          forms:
            name:
              - initial_conditions
            specific_parameters:
              - sorted_batches: 0

      - sorted_batches:
          optional: "yes"
          forms:
            name:
              - initial_conditions
            specific_parameters:
              - sorted_batches: 1

scenarios:
  - ["health_conditions", "batch_composition"]

input_forms:
  - initial_conditions:
      - risk_level:
          type: CharField
          label: "Risk level of calves"
          required: True
          help_text: "BRD risk associated with calves at purchase"
          widget: forms.Select

      - batch_isolation:
          type: CharField
          label: "Batch isolation"
          required: True
          help_text: "Level of isolation between batches"
          widget: forms.Select

      - pathogen:
          type: CharField
          label: "Main pathogen"
          required: True
          help_text: "Pathogen identified as major cause of BRD"
          widget: forms.Select

      - cost_of_antiobiotic_dose:
          type: IntegerField
          label: "Cost of antiobiotic dose"
          required: True
          help_text: "cost_of_antiobiotic_dose"
          widget: NumberInput

      - cost_purchase_calf:
          type: IntegerField
          label: "Cost of a 7 month-beef calf"
          required: True
          help_text: "Cost of a 7 month-beef calf"
```

```yaml
            widget: NumberInput

        - selling_price_bull:
            type: IntegerField
            label: "Selling price of fattened bull"
            required: True
            help_text: "selling_price_bull"
            widget: NumberInput

        - total_feeding_cost:
            type: IntegerField
            label: "Total feeding cost"
            required: True
            help_text: "total_feeding_cost"
            widget: NumberInput

        - daily_disease_impact:
            type: FloatField
            label: "Daily disease impact (€)"
            required: True
            help_text: "daily_disease_impact"
            widget: NumberInput

        - number_batches:
            type: IntegerField
            label: "Number of simultaneous batches"
            required: True
            widget: NumberInput
            widget_attrs:
              'data-min': 1
              'data-step': 1
              'data-value': 3
              'data-skin': 'round'
              'data-label': 'batches'
              'data-grid': 'true'
              'data-grid-num': 9
              'data-max': 10
        - batch_size:
            type: IntegerField
            label: "Number of calves per batch"
            required: True
            widget: NumberInput
            widget_attrs:
              'data-min': 0
              'data-step': 1
              'data-value': 10
              'data-skin': 'round'
              'data-label': 'animals per batch'
              'data-grid': 'true'
              'data-grid-num': 6
              'data-max': 30

    - execution:
        - total_duration:
            type: IntegerField
            label: "Simulation duration (half days)"
            required: True
            widget: NumberInput
            widget_attrs:
              'data-min': 0
              'data-step': 2
```

```
            'data-value': 60
            'data-skin': 'round'
            'data-label': 'half days'
            'data-grid': 'true'
            'data-grid-num': 5
            'data-max': 100
      - simulation_name:
          type: CharField
          label: "Simulation name"
          required: True
          widget: forms.TextInput

      - simulation_description:
          type: CharField
          label: "Simulation description"
          required: False
          widget: forms.Textarea

statistics:
  for_each_scenario:
    - mean_C:
        - desc: "Animals with severe clinical signs (mean, q05, q95)"
        - target_variable: [ "metapop_total_C" ]
        - groupby: [ "step" ]
        - summarise:
            - mean
            - perc_0.5:
                value: "quantile(..., 0.05)"
                desc: "5th percentile"
            - perc_0.95:
                value: "quantile(..., 0.95)"
                desc: "95th percentile"
    - nb_doses:
        - desc: "Number of doses at farm scale"
        - target_variable: [ "metapop_nb_doses" ]
        - condition: [ 'step == MAX.STEP' ]

    - nb_doses_per_batch:
        - desc: "Number of doses at batch scale"
        - target_variable: [ "batch_nb_doses" ]
        - condition: [ 'step == MAX.STEP' ]
        - add_new_column:
            batch:
              value: "as.factor(population_id + 1)"
              desc: "number of the batch"

    - mean_C_per_batch:
        - desc: "Animals with severe clinical signs (mean, q05, q95)"
        - target_variable: [ "C" ]
        - add_new_column:
            batch:
              value: "as.factor(population_id + 1)"
              desc: "number of the batch"
        - groupby: [ "step", "batch" ]
        - summarise:
            - mean
            - perc_0.5:
                value: "quantile(..., 0.05)"
                desc: "5th percentile"
            - perc_0.95:
                value: "quantile(..., 0.95)"
```

```
                desc: "95th percentile"

    - economics:
        - desc: "Net gain of fattening"
        - target_variable: ["total_population", "metapop_nb_doses",
"metapop_clinical_duration"]
        - condition: [ 'step == MAX.STEP' ]
        - add_new_column:
            gain:
              value: "selling_price_bull *  total_population -
cost_purchase_calf * initial_population - total_feeding_cost -
cost_of_antiobiotic_dose * metapop_nb_doses + daily_disease_impact *
metapop_clinical_duration / 24"
              desc: "Net gain"
  comparison_of_scenarios: []

outputs_pages:
  - Batches
  - Building
  - Economics

graphics:
  plot_mean_C:
    - data: "mean_C"
    - on_page: ["Building"]
    - label: ["Mean Clinical signs in all scenarios"]
    - single_graph: "yes"
    - scenario: ["disease_mixed_batches", "disease_sorted_batches"]
    - plot_variables:
        x: "step"
        y: "mean_metapop_total_C"
    - plot_type:
        - line:
            plot_variables:
              colour: "scenario_name"
            plot_options:
              linewidth: "2"
        - ribbon:
            plot_variables:
              ymax: "perc_0.95_metapop_total_C"
              ymin: "perc_0.5_metapop_total_C"
              fill: "scenario_name"
            plot_options:
              alpha: "0.2"
    - plot_annotations:
        title: "Animals with severe clinical signs"
        x_title: "Time (1/2 days)"
        y_title: "Average"

  plot_mean_C_disease_mixed_per_batch:
    - data: "mean_C_per_batch"
    - on_page: ["Batches"]
    - label: ["Mean Clinical per batch disease_mixed_batches"]
    - single_graph: "no"
    - scenario: ["disease_mixed_batches"]
    - plot_variables:
        group: "batch"
    - plot_type:
        - line:
            plot_variables:
              x: "step"
```

```
            y: "mean_C"
            colour: "batch"
          plot_options:
            linewidth: "2"
      - ribbon:
          plot_variables:
            x: "step"
            ymax: "perc_0.95_C"
            ymin: "perc_0.5_C"
            fill: "batch"
          plot_options:
            alpha: "0.2"
  - plot_annotations:
      title: "Animals with severe clinical signs (mixed batches)"
      x_title: "Time (1/2 days)"
      y_title: "Average"

plot_mean_C_disease_sorted_per_batch:
  - data: "mean_C_per_batch"
  - on_page: ["Batches"]
  - label: ["Mean Clinical per batch disease_sorted_batches"]
  - single_graph: "no"
  - scenario: ["disease_sorted_batches"]
  - plot_variables:
      group: "batch"
  - plot_type:
      - line:
          plot_variables:
            x: "step"
            y: "mean_C"
            colour: "batch"
          plot_options:
            linewidth: "2"
      - ribbon:
          plot_variables:
            x: "step"
            ymax: "perc_0.95_C"
            ymin: "perc_0.5_C"
            fill: "batch"
          plot_options:
            alpha: "0.2"
  - plot_annotations:
      title: "Animals with severe clinical signs (sorted batches)"
      x_title: "Time (1/2 days)"
      y_title: "Average"

plot_doses_per_batch:
  - data: "nb_doses_per_batch"
  - on_page: ["Batches"]
  - label: ["Number of doses per batch in all scenarios"]
  - single_graph: "yes"
  - scenario: ["disease_mixed_batches", "disease_sorted_batches"]
  - plot_variables:
      y: "batch_nb_doses"
  - plot_type:
      - boxplot:
          plot_variables:
            x: "scenario_name"
            fill: "batch"
  - plot_annotations:
      title: "Antimicrobial usage"
```

```
          x_title: "scenario"
          y_title: "number of doses"

    plot_doses:
      - data: "nb_doses"
      - on_page: ["Building"]
      - label: ["comparison dose in all scenarios"]
      - single_graph: "yes"
      - scenario: ["disease_mixed_batches", "disease_sorted_batches"]
      - plot_variables:
          y: "metapop_nb_doses"
      - plot_type:
          - boxplot:
              plot_variables:
                x: "scenario_name"
              plot_options:
                fill: "orange"

      - plot_annotations:
          title: "Antimicrobial usage"
          x_title: "scenario"
          y_title: "number of doses"

    plot_gains:
      - data: "economics"
      - on_page: ["Economics"]
      - label: ["Net gain"]
      - single_graph: "yes"
      - scenario: ["disease_mixed_batches", "disease_sorted_batches"]
      - plot_variables:
          y: "gain"
      - plot_type:
          - boxplot:
              plot_variables:
                x: "scenario_name"
              plot_options:
                fill: "green"
      - plot_annotations:
          title: "Net gain"
          x_title: "scenario"
          y_title: "Value"
```
